# The amyloidogenicity of the influenza virus PB1-derived peptide sheds light on its antiviral activity

**DOI:** 10.1101/211284

**Authors:** Yana A. Zabrodskaya, Dmitry V. Lebedev, Marja A. Egorova, Aram A. Shaldzhyan, Alexey V. Shvetsov, Alexander I. Kuklin, Daria S. Vinogradova, Nikolay V. Klopov, Oleg V. Matusevich, Taisiia A. Cheremnykh, Rajeev Dattani, Vladimir V. Egorov

## Abstract

The influenza virus polymerase complex is a promising target for new antiviral drug development. It is known that, within the influenza virus polymerase complex, the PB1 subunit region from the 1^st^ to the 25^th^ amino acid residues has to be is in an alpha-helical conformation for proper interaction with the PA subunit. We have previously shown that PB1(6–13) peptide at low concentrations is able to interact with the PB1 subunit N-terminal region in a peptide model which shows aggregate formation and antiviral activity *in cell cultures*.

In this paper, it was shown that PB1(6–13) peptide is prone to form the amyloid-like fibrillar aggregates. The peptide homo-oligomerization kinetics were examined, and the affinity and characteristic interaction time of PB1(6–13) peptide monomers and the influenza virus polymerase complex PB1 subunit N-terminal region were evaluated by the SPR and TR-SAXS methods. Based on the data obtained, a hypothesis about the PB1(6–13) peptide mechanism of action was proposed: the peptide in its monomeric form is capable of altering the conformation of the PB1 subunit N-terminal region, causing a change from an alpha helix to a beta structure. This conformational change disrupts PB1 and PA subunit interaction and, by that mechanism, the peptide displays antiviral activity.

**Graphical abstract:** 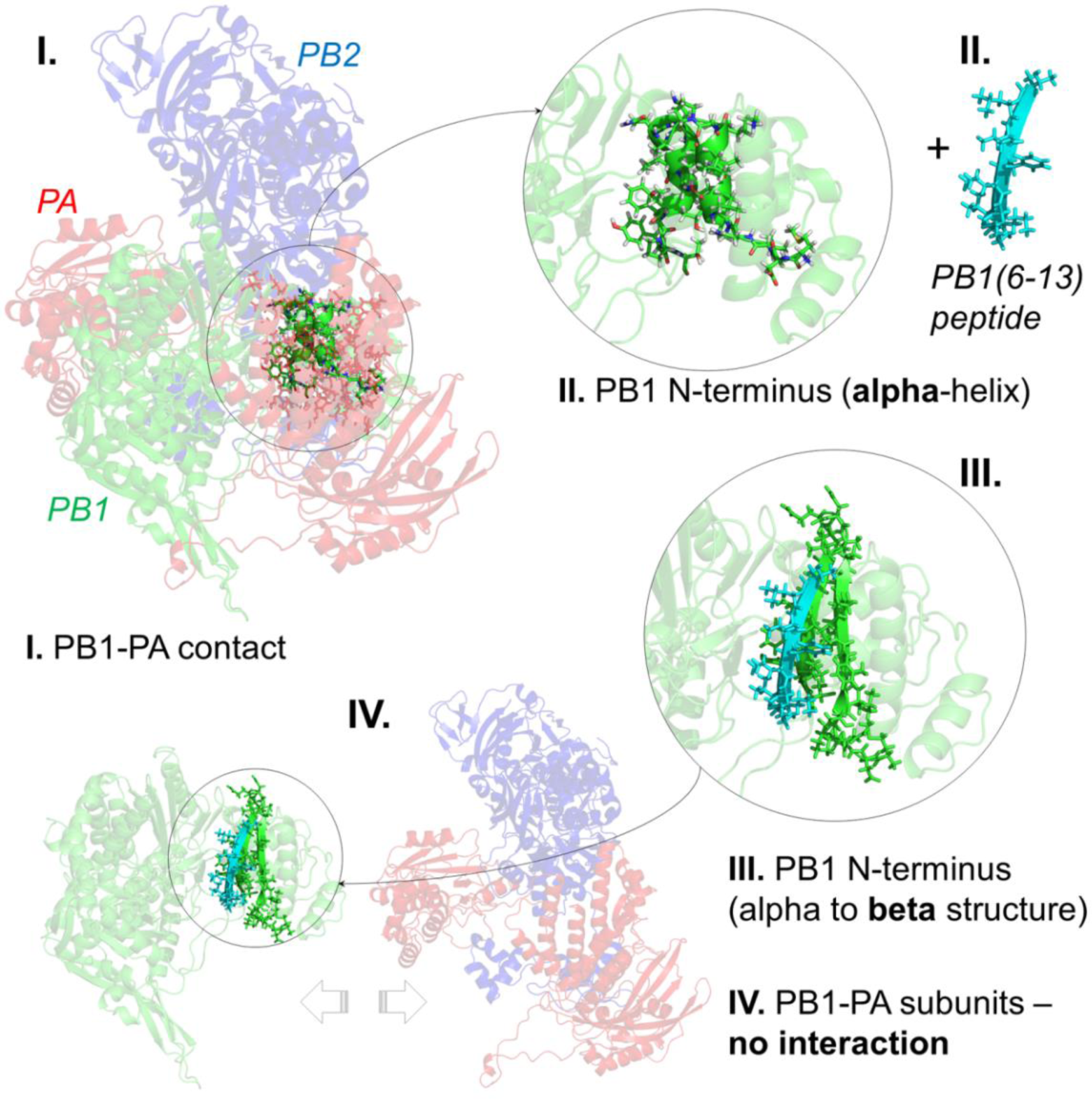

## Introduction

At the moment, there are a very limited number of anti-influenza drugs. Neuraminidase and the M2 ion channel are the most often targeted viral proteins in antiviral therapy [1]. However, due to the wide variability in influenza viruses (through mutation), resistance to existing drugs has appeared in some strains. This leads both to the necessity for new drugs for pre-existing molecular targets, and to the need to search for new targets [2]. There are some antiviral drugs which affect influenza virus hemagglutinin [3]. Currently, the influenza virus polymerase complex [4], [5] and nucleoprotein [6]–[9], which have relatively conservative amino acid sequences, are considered as the most promising targets for drug development.

The polymerase complex consists of three subunits: PA, PB1, and PB2. An increased interest in research and development devoted to the polymerase-complex-affecting drugs occurred after the publication of the polymerase complex’s structure (obtained by means of X-ray diffraction analysis and cryoelectron microscopy) [10]–[12]. It was shown that a site, from the 1^st^ to the 15^th^ amino acid residues of the influenza virus polymerase complex PB1 subunit (hereinafter PB1(1–15)), is located at a PB1/PA subunit interface which plays a key role in the process of their interaction, and is in the conformation of an alpha helix [10], [11]. It should be noted that, when delivered into cells, the PB1(1–25) and PB1(1–15) peptides have antiviral activity against the influenza A virus [13].

We have previously shown that the PB1(6–13) fragment is able to interact with the PB1 subunit N-terminal region in a peptide model [14] and has antiviral activity *in vitro*[15].

## Methods

### Peptides

Peptides used as a model for the influenza A virus polymerase complex PB1 subunit N-terminal region: MDVNPTLLFLKVPAQNAISTTFPYT (PB1(1–25)) and TLLFLKVPAQNAISTTFPYT (PB1 (6–25)). The peptides tested were: TLLFLKVP (PB1(6–13)); TLLFLKVPA (PB1(6–13)-Ala); and FITC-labeled TLLFLKVPA (FITC-PB1(6–13)-Ala). All peptides were synthesized at the Saint-Petersburg State University as described in referenced work [15].

### Preparation of peptides in different aggregation states

Evaluation of the PB1(6–13)/PB1(6–13)-Ala peptide aggregation during dissolution was carried out as follows. The dissolved peptide was centrifuged at 20,000 g, 4°C, for 40 minutes. The supernatant was collected and mixed with acetonitrile (AN) to a final concentration of 40%, and reverse phase chromatography was performed: column-YMC-Triart C18, 2.0×150 mm, granule size 3 μm; mobile phase A – 0.1% TFA (Sigma, USA) in deionized water, mobile phase B – 0.1% TFA in AN, isocratic mode, composition A: B – 68:32, flow rate 0.2 ml/min, detection by optical density at 214 nm, column temperature 25°C, loop volume 100 μl, sample volume 5 μl, assay time 25 minutes.

Peaks corresponding to the peptide yield time (retention time, Rt, was 12.6 min ± 0.1 min) were integrated using the Waters chromatograph software (USA). In order to determine the concentration of the peptide in soluble form remaining in the supernatant after centrifugation, peptide peak areas in the solvent and in AN, at the same concentrations, were compared, assuming the peptide solubility in 40% AN to be equal to 100%.

If the peptide concentration in the supernatant corresponded to the peptide concentration in AN, and a right shift in the Congo red absorption spectrum (indicative of beta-structured aggregates) was absent, the state was termed “monomers”. The state was termed “fibrils” in the case of precipitate formation after centrifugation (containing amyloid-like fibrils according to atomic force microscopy), accompanied by Congo red dye-binding assay spectral right shift.

### Congo red dye (CR)

Congo red dye (Sigma Aldrich, USA) at a concentration of 50 μM in PBS was mixed with the 10 μM peptide. As a control, a similar sample was used in which a buffer was added instead of the peptide. Absorption spectra were recorded on an Avantes Ava Spec 2048 instrument (Avantes, The Netherlands). The Congo red dye absorption spectrum characteristic maximum is in the region of 500 nm (the exact value depends on the buffer used). In the presence of beta structures in the sample, the dye binds to them, which leads to a shift of its characteristic maximum to the right by 10–20 nm. The spectra were visualized using Origin2015 software.

### Atomic force microscopy (AFM)

Five microliters of PB1(6–13) peptide, at a concentration of 1 mM in PBS, was applied to a freshly cleaved mica surface (SPI supplies, USA) and incubated for about 30 seconds, after which the sample was washed three times with 50 μl of water and dried in a thermostat for 30 minutes at 55°C. The scanning was performed in a semi-contact mode on the SolverNext atomic force microscope (NT-MDT, Russia, Zelenograd) under the NovaPx program (production of NT-MDT) using an NSG-03 probe. The resulting images were processed in the Gwyddion software [16].

### Small-angle neutron scattering (SANS)

In order to measure the peptides’ scattering spectra by the SANS method, solutions were prepared in PBS/D_2_O at a concentration of 1 mM. The spectra were obtained with the YuMO spectrometer located on the fourth channel of the IBR-2 high-flux reactor (Frank Laboratory of Neutron Physics, Joint Institute for Nuclear Research, Dubna, Russia). The measurements were carried out in the standard geometry described previously [17], [18]. The scattering detection was performed by two detectors simultaneously in the Q region from 0.006 to 0.3 Å. Preliminary processing of the data was also carried out according to the scheme described in the reference [19]. In the method, obtaining and normalizing the curves, in absolute units, is realized with the help of metallic vanadium located directly before the detector. The curves were visualized using Origin2015 software.

### Surface plasmon resonance (SPR)

Activation of the CM5 sensor chip (GE Healthcare, USA) and immobilization of PB1(6–13)-Ala peptide monomer on its surface were performed according to the manufacturer's instruction (GE Healthcare, USA) on the Biacore X100 device (GE Healthcare, USA). Briefly, after chip surface activation, a ligand solution (at a concentration of 400 μM in MES (Sigma, USA), pH 6.0, 2% ethanol) was injected at a flow rate of 10 μl/min, contact time of 900 s, after which, the unoccupied channel was inactivated with ethanolamine, pH 8.0. The level of immobilization was 3100 RU (1 RU = 1 pg/mm^2^). The same procedures, but without the addition of peptide, were performed with a reference channel.

PB1(6–13)-Ala and PB1 (6–25) peptide monomers, dissolved into five different final concentrations (59 μM, 89 μM, 133 μM, 200 μM, and 300 μM) in HEPES with 2% ethanol, were used to measure K_d_ using a flow rate of 30 μl/min, a contact time of 180 s, and a dissociation time of 180 s. The measurements were carried out in the single cycle mode. Sensograms were processed in the Biacore software and were visualized using Origin2015 software.

### Time-resolved small-angle X-ray scattering (TR-SAXS)

TR-SAXS experiments were carried out at the synchrotron radiation source ESRF (Grenoble, France) using the ID02 spectrometer in accordance with the corresponding protocols. Peptide monomers were dissolved in PBS with 10% DMSO and then mixed in an equimolar ratio, at final concentrations of 0.2 mM, using a “stopped flow” system. Five milliseconds after mixing, the 100 SAXS spectra registration was measured at 3 s intervals.

Data processing was performed using original software developed by N.V. Klopov et al., NRC KI PNPI, Gatchina, Russia. The program is built around the singular value decomposition (SVD) method and is capable of identifying the main spectral components (U(q)) and the variation of such components over time (V(t)). Based on the exponential kinetics of the main components, the SAXS spectra for the initial (t=0) and the final (t=∞) states of the system were calculated in order to determine the state of the peptide system before and after mixing. Spectra were visualized using Origin2015 software.

### MicroScale Thermophoresis (MST)

Experiments using MST were performed on the Monolith NT.115 (NanoTemper Technologies GmbH, Germany), in accordance with manufacturer protocols using 80% laser intensity, 25^o^C. A 0.5 μM concentration of the FITC-PB1(6–13)-Ala peptide and a range of 16 PB1(1–25) peptide concentrations (from 0.012 to 400 μM) were used. The measurements were carried out in PBS containing 10% DMSO. In the concentrations used, the peptides were in the form of monomers. The data were processed in instrument software and were visualized using Origin2015 software.

## Results

### PB1(6–13) peptide forms amyloid-like fibrils in PBS

It was previously shown that PB1(6–13) forms aggregates in high (greater than 1 mM) concentrations in PBS. According to reversed-phase chromatography data (not shown), it was found that almost all of the peptide aggregates in these conditions, and only a small part of it remains in the form of monomers. Figure 1 shows the resulting PB1(6–13) peptide aggregates using atomic force microscopy.

**Figure 1.**
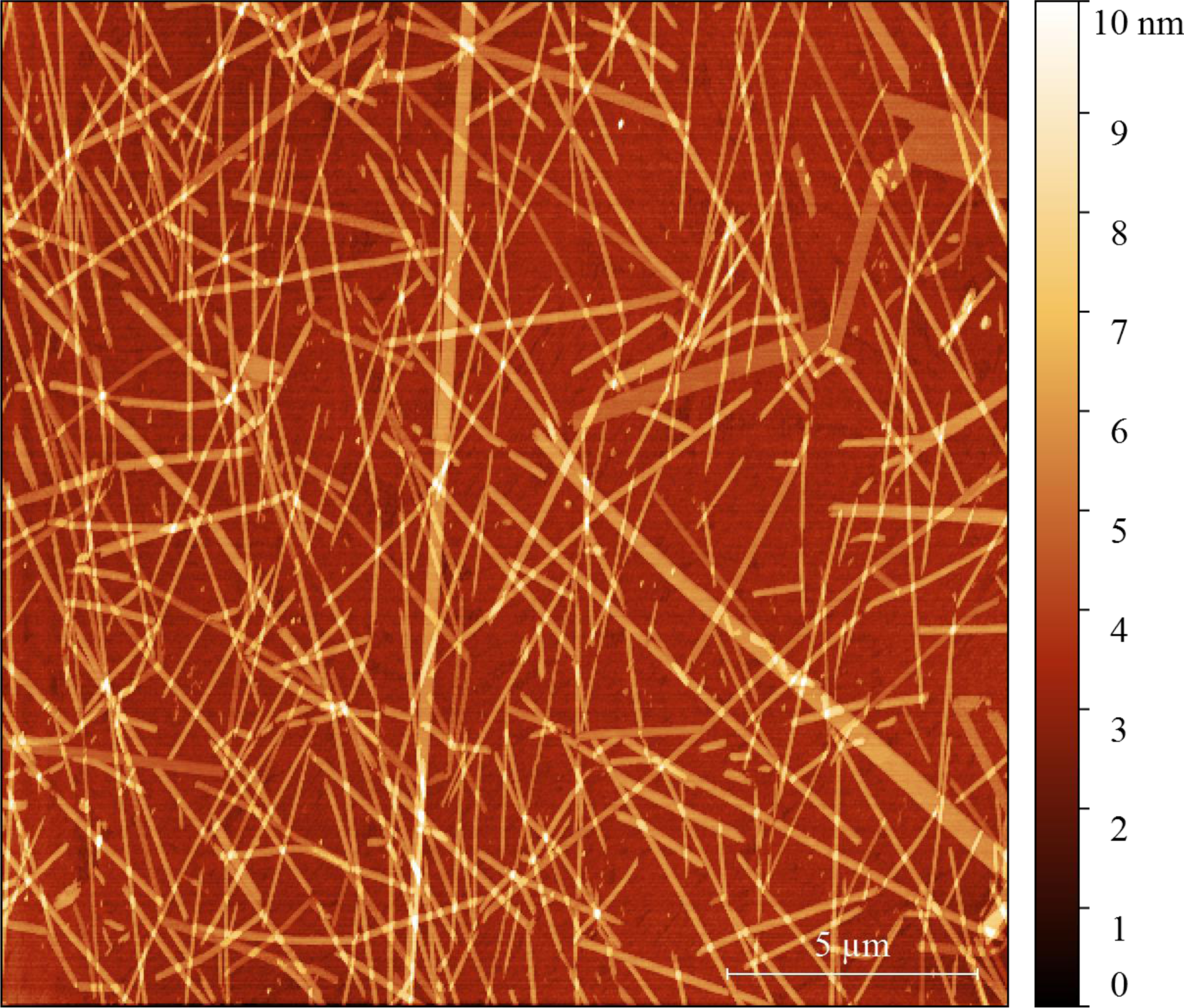
Atomic force microscopy of aggregates formed upon PB1(6–13) peptide dissolution at a concentration of 1 mM in PBS

The morphology of the aggregates seen is similar to that of amyloid-like fibrils.

Congo red, a specific dye whose spectrum shifts to the right when bound to amyloid-like fibrils, was used for this aggregate binding test [20]. The absorbance spectra of Congo red dye and peptide sample precipitate mixtures are shown in Figure 2.

**Figure 2.**
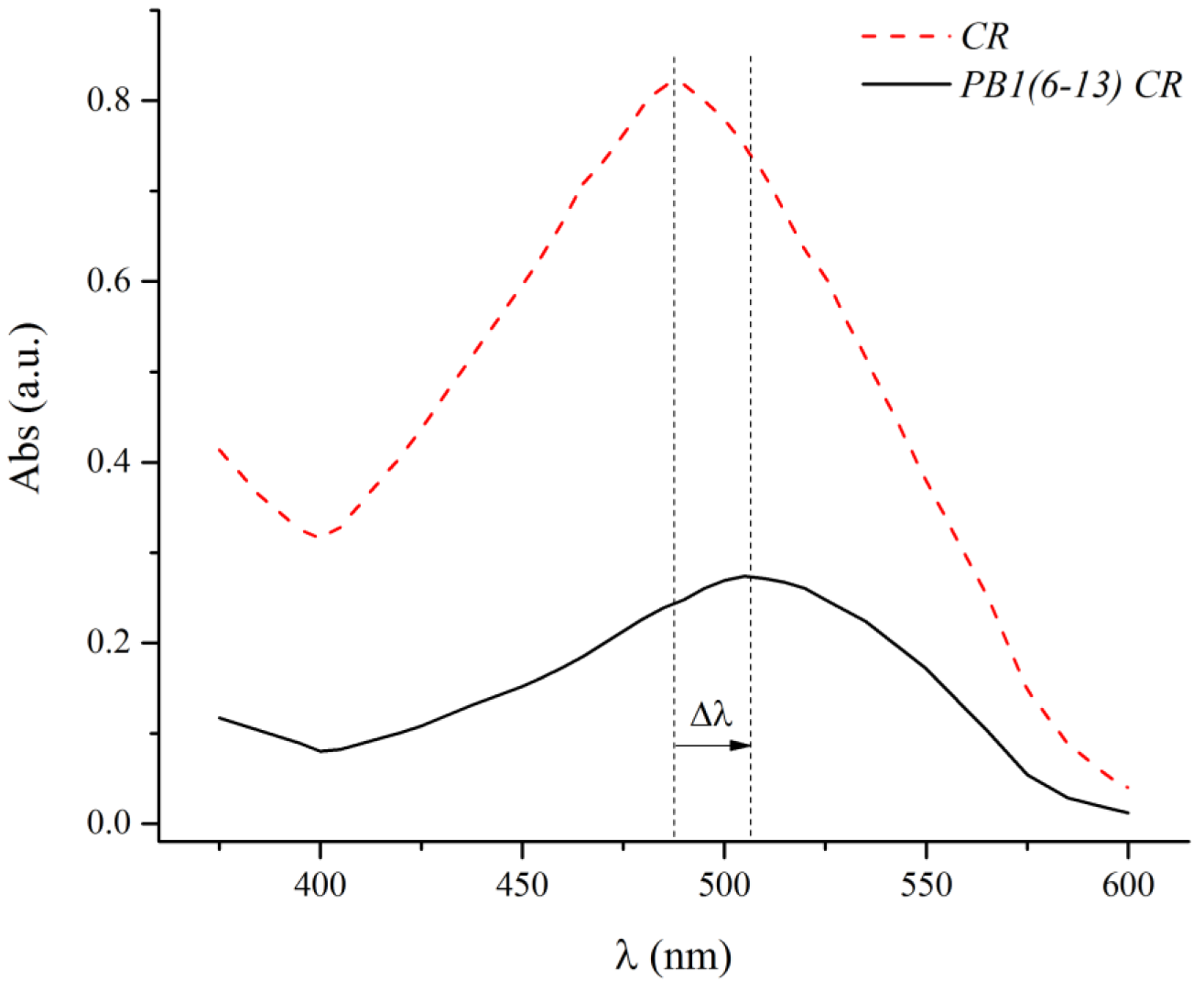
The PB1(6–13) peptide absorption spectrum in the presence of Congo red is indicated by a black solid curve; the Congo red spectrum alone is indicated by a red dashed curve. In the presence of the peptide, a characteristic Congo red absorption maximum right shift (Δλ) is observed

Thus, it was shown that the PB1(6–13) peptide forms amyloid-like fibrils upon dissolution in PBS. Small-angle neutron scattering (SANS) data also indicates that the sample contains not individual peptide molecules, but aggregates (Figure 3). The initial segment of the SANS spectrum followed a power law function, indicating that in solution the protofibrils formed by the peptide do not have an extended rod-like shape but instead exhibit the properties of a polymer chain in poor solvent.

**Figure 3.**
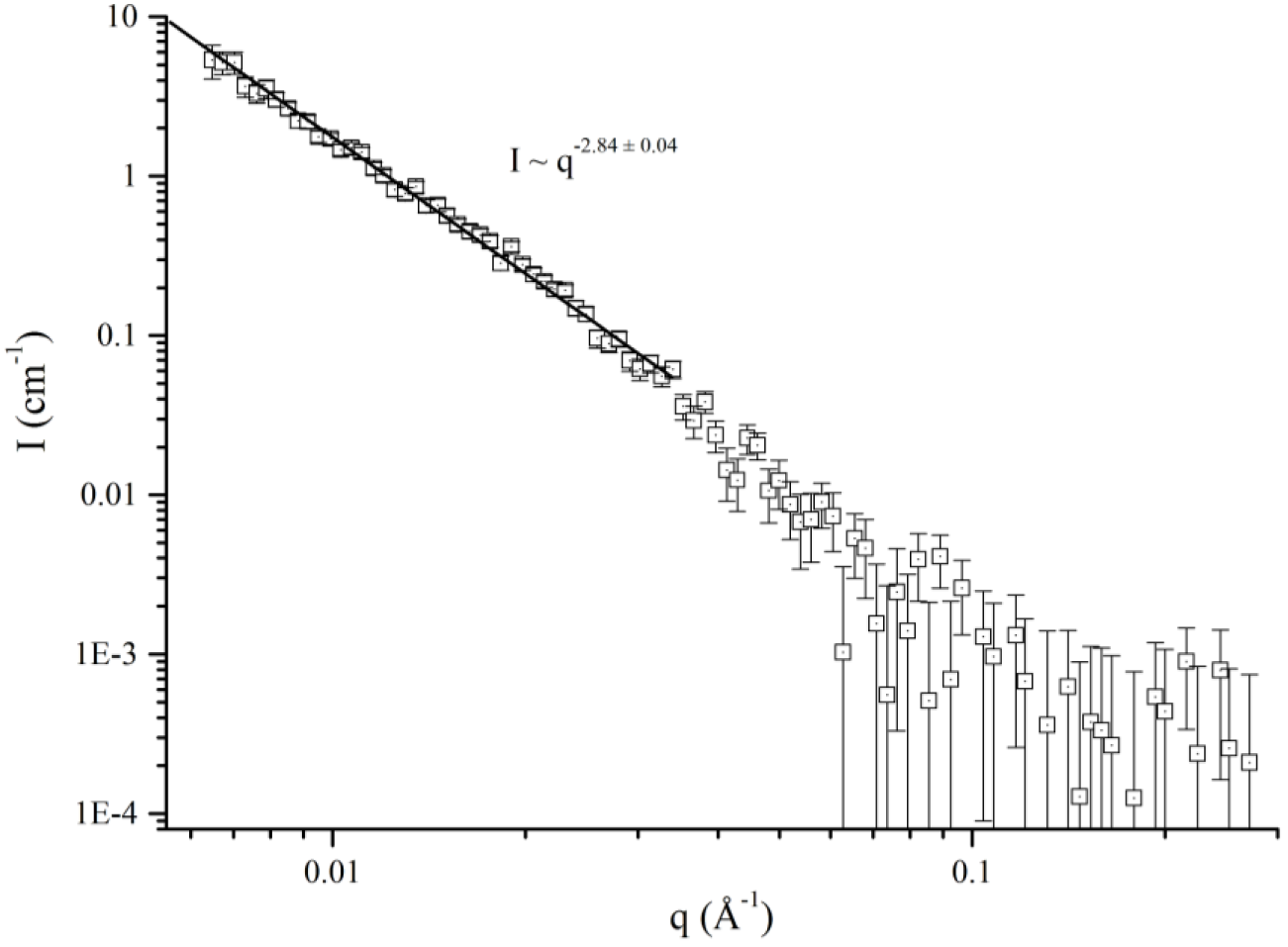
PB1(6–13) peptide SANS spectra in PBS/D_2_O. The data are approximated by a power law using the least squares method. The power law exponent of 2.84 ± 0.04 corresponds to chain in poor solvent

### Kinetics of fibril formation

Monomers of PB1(6–13)-Ala peptide were immobilized on the chip surface in order to study the kinetics of fibril formation by surface plasmon resonance (SPR) in a manner similar to previous work [21]. Five different concentrations (59, 89, 133, 200, and 300 μM) of the same peptide monomers, dissolved in HEPES containing 2% ethanol, were used as the analyte (Figure 4).

**Figure 4.**
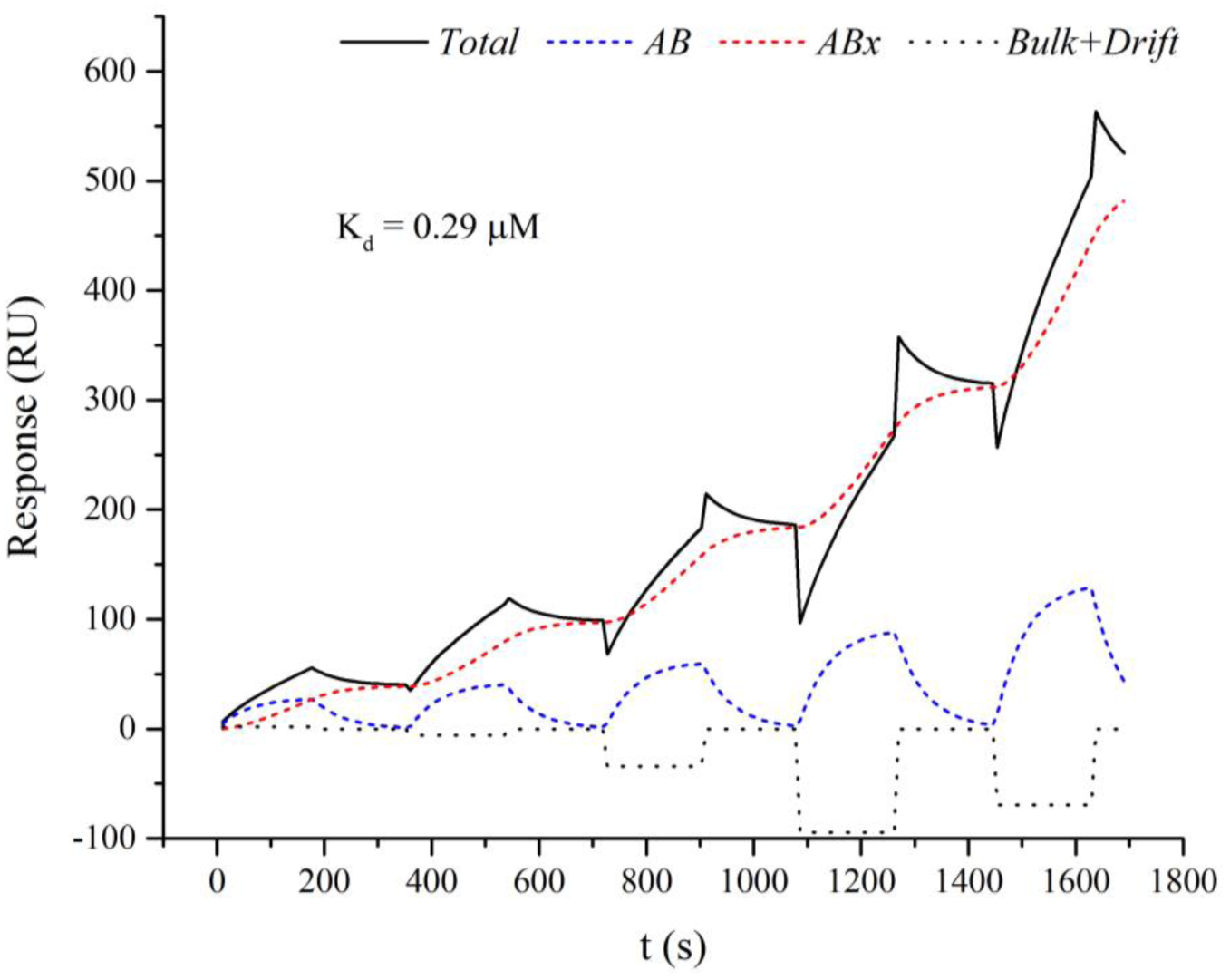
Sensogram of: the PB1(6–13)-Ala peptide interaction with itself (black solid curve); approximated by a two-step reaction model (blue and red curves); and corrected for bulk and drift effects (black dotted curve)

This interaction cannot be described by the Langmuir kinetics, but it is well described by a two-step reaction model with correction for bulk and drift effects. The equilibrium dissociation constant (K_d_) calculated from this approximation was 0.29 μM, which indicates the propensity of the peptide to homo-oligomerization. The data obtained also point to the tendency of the peptide to form fibrils according to the mechanism described for amyloidogenic polypeptides.

### The PB1(6–13) peptide fibril’s ability to interact with the PB1 polymerase subunit N-terminal region in the peptide model

A study of the PB1(6–13) peptide’s interaction with the influenza A virus PB1 polymerase subunit was carried out in a peptide model in which the PB1 N-terminal region (6^th^ to 25^th^amino acid residues) plays the role of the PB1 polymerase subunit. According to the literature, this fragment of the influenza A virus PB1 subunit contains key amino acid residues necessary for interaction with the PA subunit [22]. At the first stage, the interaction between the PB1(6–13) peptide fibrils and the PB1 subunit N-terminal region was studied by the SANS method (Figure 5).

**Figure 5.**
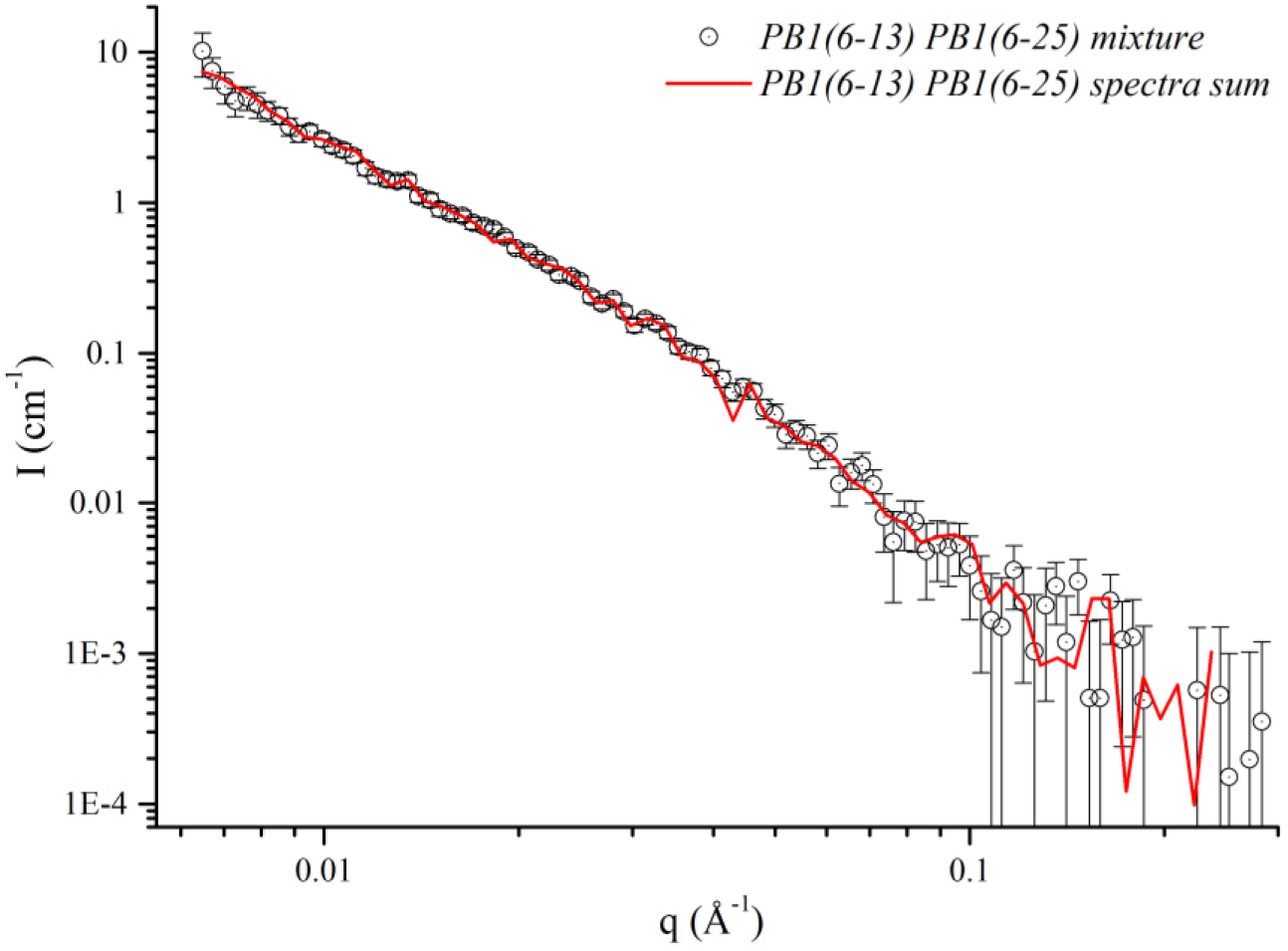
SANS spectra of: a peptide mixture solution containing PB1(6–25) and PB1(6–13) in fibrillar form (black dots); and the normalized mathematical sum of the individual scattering curves for PB1(6–25) and PB1(6–13) (red curve)

When analyzing the SANS data, it was found that the spectrum of the peptide mixture (black dots) corresponds to the mathematical sum of the individual component’s spectra (red curve). It can be concluded that after the fibril’s formation, the peptides are no longer capable of interacting, and that the active form of the peptide PB1(6–13) is, most likely, the monomer. Further, the interaction parameters were studied using the PB1(6–13) peptide monomer.

### Timing of the interaction between the PB1(6–13) peptide and the PB1 subunit N-terminal region

The features of the PB1(6–13) and PB1(6–25) interaction as monomers was studied by the SAXS method. This method allows not only qualitatively establishing the presence or absence of interaction, and describing the structures obtained, but also evaluating the characteristic interaction times. A set of one hundred SAXS spectra, obtained during the first five minutes after peptide mixing, was processed by the SVD method in order to identify the most variable singular decomposition spectrum components.

The variation shape and dynamics of such decomposition component in the q measurement range from 0.1 to 1.0 nm^−1^ in a PB1(6–13) and PB1(6–25) peptide mixture solution is shown in Figure 6(A). The restored initial and final states of the system are shown in Figure 6(B).

**Figure 6.**
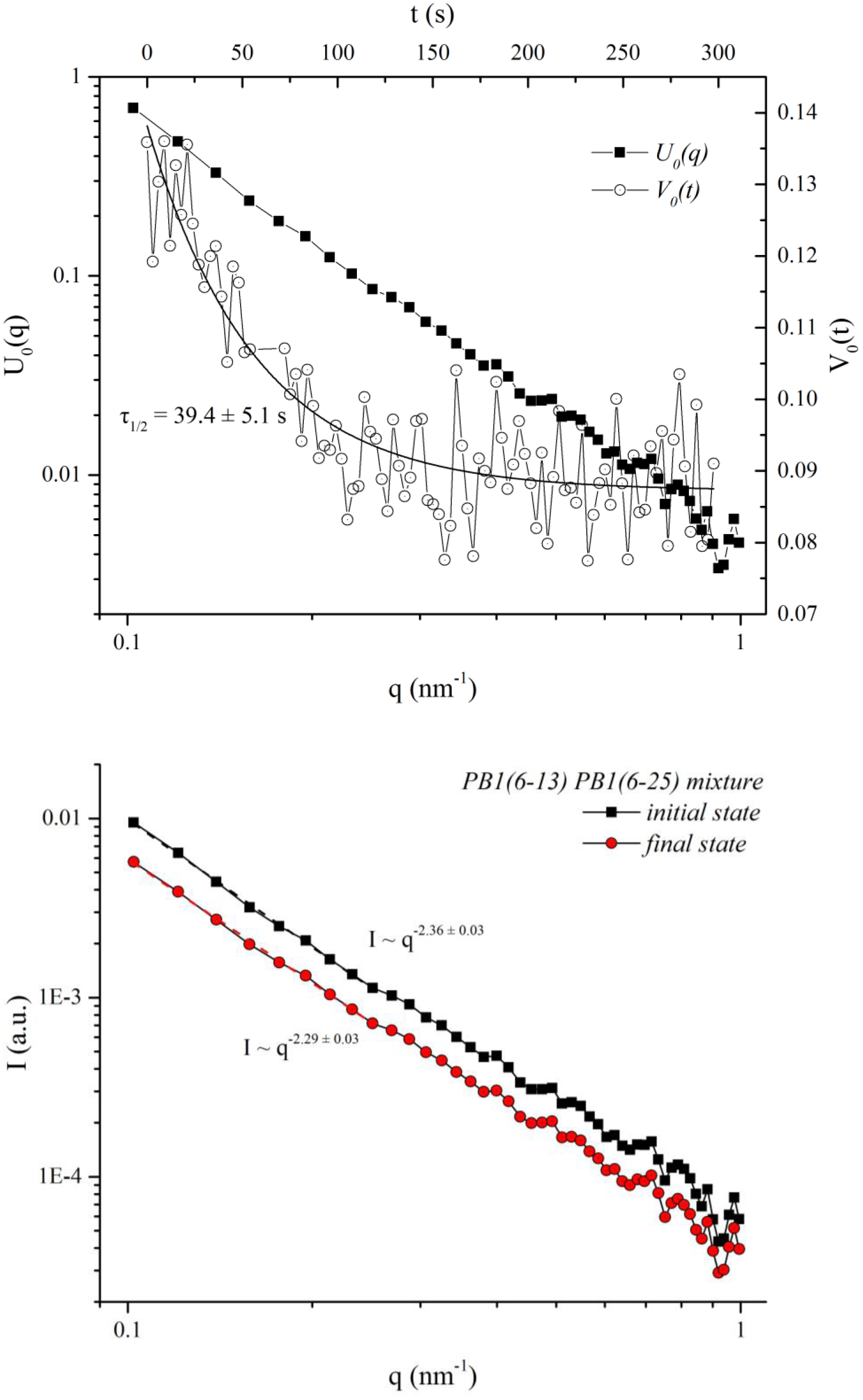
(A) (top) – U_0_(q) is the form of the SAXS spectra singular decomposition most variable (zero) component in the measurement range q from 0.1 to 1.0 nm^−1^ in the PB1(6–13) and PB1(6–25) peptide mixture solution; V_0_(t) is the change in the zero component as a time function. The characteristic time (τ_1/2_, the half-reaction time) was 39.4 ± 5.1 s. (B) (bottom) – The PB1(6–13) and PB1(6–25) peptide mixture system initial (t = 0) and final (t = ∞) states spectra, reconstructed on the basis of a change in the singular decomposition zero component

The change in the SVD zero component appears to be due to a decrease in the curves intensity; it may indicate rapid aggregation of the peptide mixture. It should be noted that there are no such spectra changes in the peptides separately (data not shown).

Analogous calculations were made in the study of the PB1(6–13) and PB1(6–25) peptides’ SAXS spectra in the q interval from 0.01 to 0.2 nm^−1^. The changes in the singular decomposition zero and first components are shown in Figure 7(A) and 7(B), respectively. The initial and final states of the system are shown in Figure 7(C).

**Figure 7.**
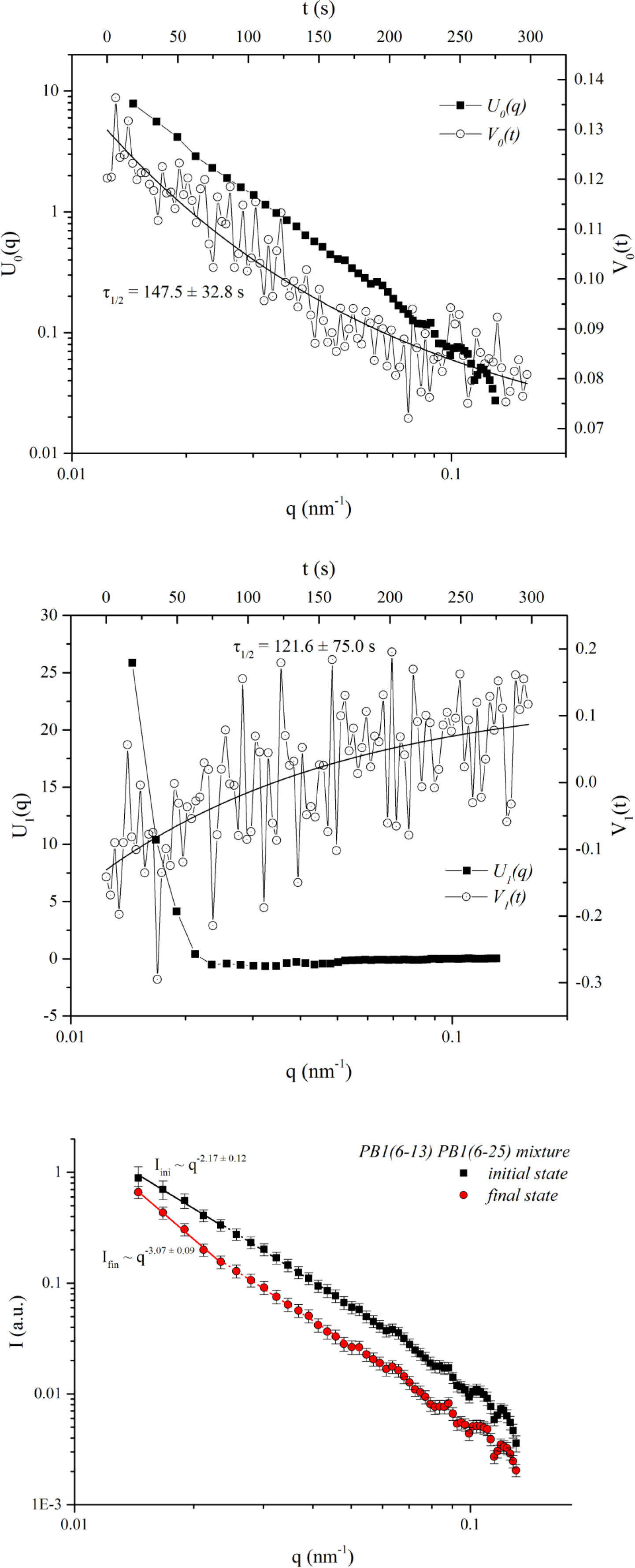
(A) (top) – U_0_(q) is the form of the SAXS spectra singular decomposition zero component in the measurement range q from 0.01 to 0.2 nm^−1^ in the solution of the PB1(6–13) and PB1(6–25) peptide mixture; V_0_(t) is the change in the zero component as a time function. The characteristic time (τ_1/2_, the half-reaction time) was 147.5 ± 32.8 s. (B) (middle) – U_1_(q) is the form of the SAXS spectra singular decomposition first component in the measurement range q from 0.01 to 0.2 nm^−1^ in the solution of the PB1(6–13) and PB1(6–25) peptide mixture; V_1_(t) is the change in the first component as a time function. The characteristic time (τ_1/2_, the half-reaction time) was 121.6 ± 75.0 s. (C) (bottom) – The PB1(6–13) and PB1(6–25) peptide mixture system initial (t = 0) and final (t = ∞) states spectra, reconstructed on the basis of a change in the singular decomposition zero and first components

In this case (Figure 7), the change in the zero component can also be attributed to a decrease in the overall SAXS spectra intensity. However, the change in the first component seems to reflect an increase in the slope of the curve’s initial section; the slope changes over a characteristic time on the order of 120s. The increase in scattering at q below 0.025 nm^−1^ can be interpreted as an effect caused by the appearance of large aggregates in the sample as a result of the peptide interactions.

### Nature of the aggregates

In order to confirm that the PB1(6–13) peptide in monomeric form is capable of inducing a conformational transition in the influenza A virus polymerase PB1 subunit N-terminal region, the monomers of PB1(6–13) and PB1(6–25) peptides, and their the mixture, were tested for the presence of beta-structured aggregates using a specific Congo red dye (Figure 8). The peptides were dissolved in 10% DMSO.

**Figure 8.**
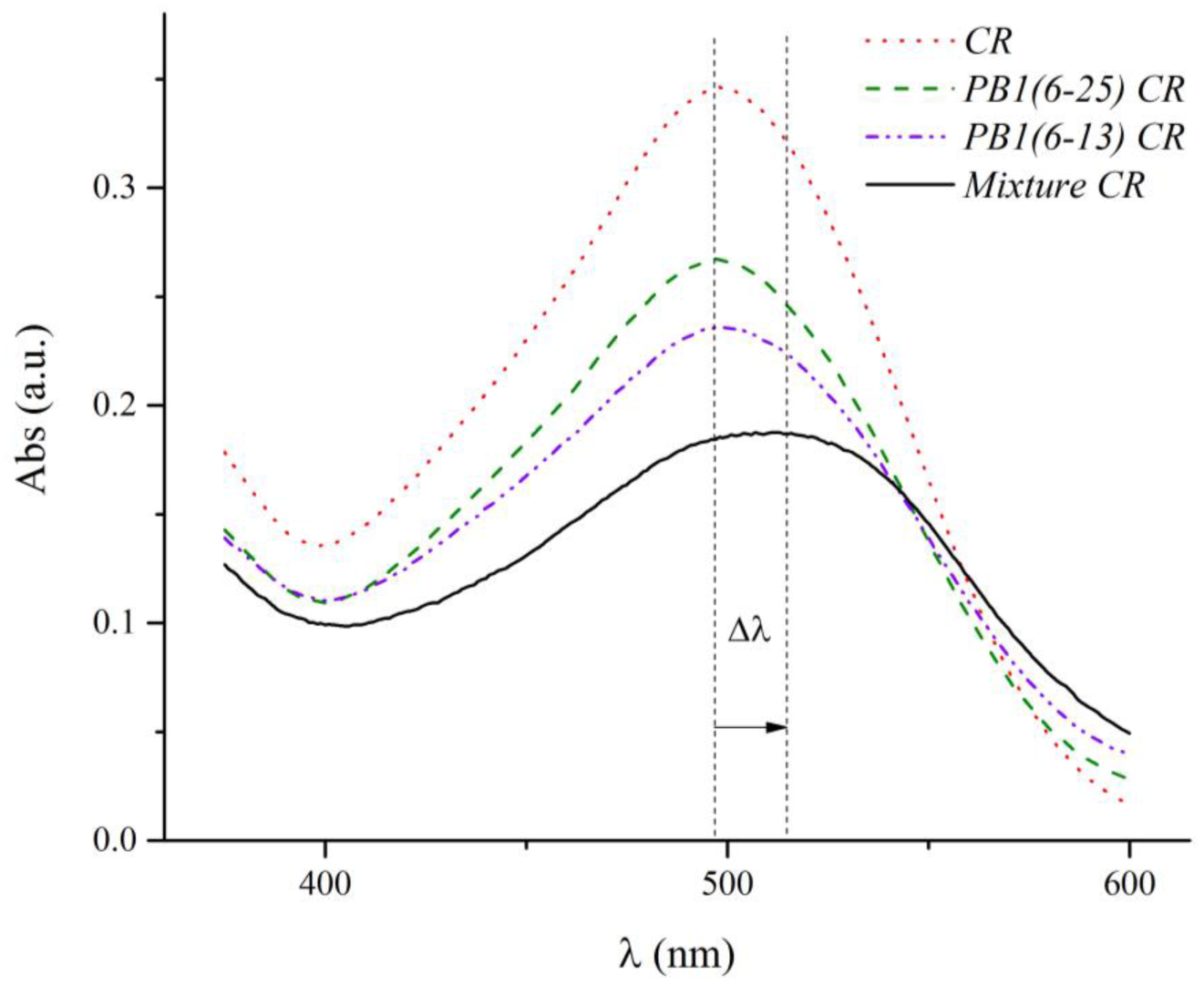
PB1(6–13) peptide monomer absorption spectrum in the presence of Congo red (blue curve); initial Congo red spectrum (red curve); PB1(6–25) peptide monomer absorption spectrum in the presence of Congo red (green curve); peptide mixture in the presence of Congo red (black curve). In the peptide mixture, the Congo red absorption maximum right shift (Δλ), which is characteristic for beta-structured aggregates, is observed

The aggregates formed as a result of the peptides’ interaction are of an amyloid-like nature. The peptides’ starting solutions in the buffer containing DMSO do not bind CR. Thus, the PB1(6–13) peptide monomer is able to induce a conformational transition in the influenza A virus polymerase PB1 subunit N-terminal region in the peptide model.

### PB1(6–13) peptide monomer and PB1 subunit N-terminus equilibrium dissociation constant

The equilibrium dissociation constant for the interaction between PB1(6–13) peptide monomers and PB1 subunit N-terminal regions (from 6^th^ to 25^th^ amino acid residues) was determined by surface plasmon resonance and MicroScale Thermophoresis (MST). The sensogram obtained by SPR is shown in Figure 9. The PB1(6–13)-Ala peptide was immobilized on the CM5 chip as the ligand, and the PB1(6–25) peptide in various concentrations (59, 89, 133, 200, and 300 μM), in HEPES buffer containing 2% ethanol, was used as analyte.

**Figure 9.**
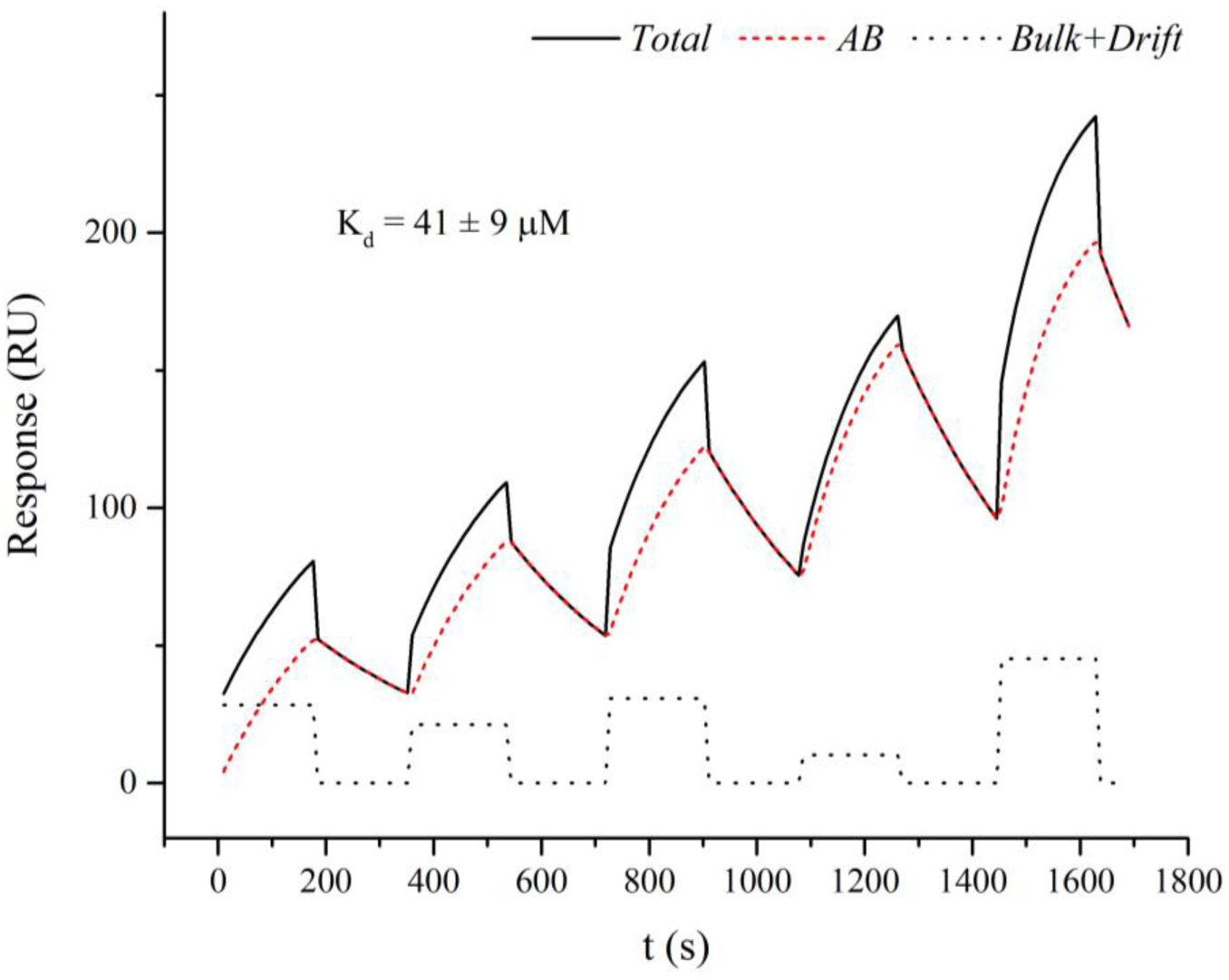
Sensogram of: the PB1(6–13)-Ala peptide monomer interaction with PB1(6–25) in solution (black solid curve); approximated by a Langmuir kinetic equation (red curve); and corrected for bulk and drift effects (red dotted curve)

In contrast to the interaction of PB1(6–13)-Ala peptide with itself (Figure 4), the time course of PB1(6–13)-Ala / PB1(6–25) interaction could be approximated by a Langmuir kinetic equation that implies a simple equimolar association of the two peptides. The equilibrium dissociation constant for PB1(6–13)-Ala / PB1(6–25) interaction, as determined by SPR, was 41 ± 9 μM. This value is two orders of magnitude higher than for PB1(6–13)-Ala / PB1(6–13)-Ala interaction during fibril formation.

In order to evaluate the dissociation constant for peptide interaction in bulk solution, an MST experiment was performed. In this experiment, FITC-labeled PB1(6–13)-Ala peptide was used as the test substance. The N-terminal region of the PB1 subunit was represented by the longer PB1(1–25) peptide. The test peptide was used in one concentration (0.5 μM), and the N-terminal region was mixed with it in 16 different concentrations. The relative fluorescence intensity of the FITC-PB1(6–13)-Ala peptide and PB1(1–25) peptide mixture (with various concentrations of the latter) obtained in solution by thermophoresis is shown in Figure 10.

**Figure 10.**
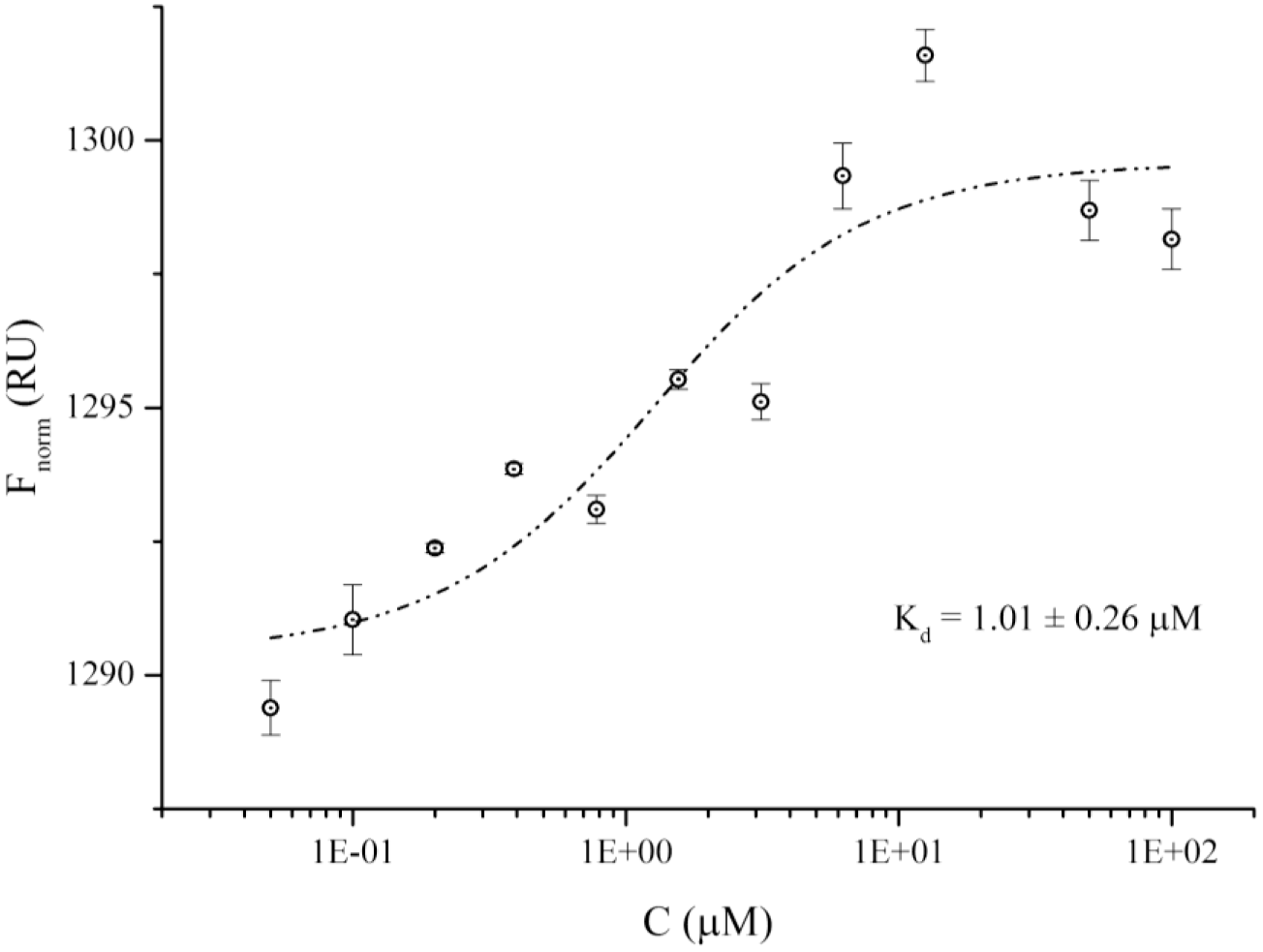
Dependence of the peptide mixture’s relative fluorescence FITC-PB1(6–13)-Ala on the PB1 subunit N-terminal region PB1(1–25) concentration, obtained by the MST (points represent the average of 3 measurements corrected for intensity drift)

In MST experiments, the equilibrium dissociation constant for the peptides’ interaction was 1.01 ±

0.26 μM. The large difference in the interaction affinities obtained by SPR and MST methods could be attributed to the effect of immobilization on the peptide conformation and the influence of the chip surface on peptide-peptide interaction, although one cannot exclude that the extra residues in PB1 N-terminal region or of the fluorescent label affect the binding affinity. It should also be noted that SPR method is ordinarily used to determine K_d_ of antigen-antibody interaction whose molecular mass is usually much higher than that of the peptide monomers [23], [24]. The low mass of peptides causes small response that could make SPR inaccurate for peptide interaction measurement. The value for the dissociation constant obtained by MicroScale Thermophoresis should therefore be considered a more reliable estimate with respect to the interaction between the PB1(6–13) peptide monomer and the PB1 subunit N-terminal region in bulk solution.

## Discussion

Previously, we showed that the peptide from the PB1 subunit N-terminal region, corresponding to 6^th^ to 13^th^ residues, has antiviral activity against the influenza A virus [15]. Some published reports have assumed that the peptide PB1(1–25), which has antiviral activity upon delivery to cells, interacts with the PA subunit at its binding region with PB1 and disrupts the polymerase complex [25]. In other words, the peptide is inserted into the PA subunit pocket, instead of the PB1 N-terminal region, thereby preventing the assembly of the polymerase complex. However, in this work, we show by SANS, AFM, CR, and SPR that the PB1(6–13) peptide, upon dissolution in phosphate buffer, forms amyloid-like fibrils with a K_d_ on the order of 0.1 μM. This fact allows us to consider another possible mechanism of action for such peptides.

The possibility of conformational transition induction in proteins, when interacting with peptides, is discussed in a review [26]. We considered, earlier, the hypothesis about these mechanisms as they relate to the interaction between the PB1(6–13) peptide and the PB1 subunit N-terminal region (6^th^ to 25^th^ residues) [14]. However, the observed phenomenon was not considered to be amyloid-like conformation transition. In light of the data obtained in this study, it can be assumed that the amyloid-like fibril-prone PB1(6–13) peptide interacts with the influenza A virus polymerase complex PB1 subunit N-terminal region (with K_d_ on the order of 1 μM, according to solution MST data), causing a conformation shift towards beta structures. *In silico* and *in vitro* experiments have shown [14] that the PB1(6–25) N-terminal region is capable of assuming a beta-conformation. In this study, via the SAXS and CR dye binding methods, it was shown that the interaction between the PB1(6–13) peptide monomer and the PB1 N-terminal region leads to the formation of large beta-structured aggregates with a characteristic time on the order of 120 s. The results obtained are in complete agreement with our previous work [14] wherein the formation of large aggregates (around 500 nm) was shown, for low concentrations of the peptides, by electron microscopy and dynamic light scattering; an increase in anti-parallel beta structure content in the mixtures was measured by the circle dichroism method.

It should be noted that after the formation of amyloid-like fibrils by the PB1(6–13) peptide itself, it is no longer able to interact with the PB1(6–25) N-terminal region, as shown in the SANS experiments. Thus, the peptide must be in a monomeric form to induce a conformational transition in the N-terminal region of the influenza PB1 subunit.

The data obtained suggest that, under the influence of the PB1(6–13) or PB1(6–13)-Ala peptide monomeric forms, the influenza A virus polymerase complex PB1 subunit N-terminal region acquires a beta conformation and loses its ability to interact with the PA subunit, since its alpha conformation is critical for such interaction[10], [11]. As a result, the influenza virus polymerase complex functionality is broken; this explains the PB1(6–13) peptide antiviral activity, shown in [15]. Several questions remain unanswered [15], such as: At what stage of the influenza virus life cycle does the peptide exerts its antiviral effect? Can it provoke disassociation of previously formed polymerase complexes? Does it affect only newly synthesized PB1subunits?

It should be noted that, in the case of peptide-mediated conformational transition induction, the study peptide itself is not consumed and it can repeatedly affect the PB1 subunit N-terminal regions. Thus, when using the peptide as an antiviral agent, much smaller quantities of the drug are required than when implementing other mechanisms proposed [25] for PB1(1–25), in which an equimolar ratio of the peptide to the PA subunit is required.

## Conclusion

In this study, it was shown that the PB1(6–13) peptide is able to interact with the influenza A virus polymerase complex PB1 subunit in a peptide model. PB1(6–13) molecules tend to form amyloid-like fibrils. Mixing of PB1(6–13) and PB1(6–25) monomeric peptides results in the formation of beta-structured aggregates which may indicate an induction of PB1 subunit N-terminus conformational transition under the influence of PB1(6–13). This peptide appears to be a promising candidate for further drug development.

## Acknowledgments

This work was supported by State contract *№*14.N08.11.0080 “Human influenza A virus antiviral peptide preclinical study” (Federal Target Program: “The Russian Federation pharmaceutical and medical industry development for the period until 2020 and beyond”); Experiment #LS2508 @ID02, ESRF, Grenoble, France (TR-SAXS); RFBR grant №14-24-01103 ofi_m “The method for structural and dynamical diagnostics of multimolecular nucleoprotein complexes by verification of the molecular dynamics models via small-angle X-ray and neutron scattering”. SANS experiments were performed at JINR, Dubna, Russia. The results of the work were obtained using computational resources of Peter the Great Sainte-Petersburg Polytechnic University Supercomputing Center (www.spbstu.ru). MicroScale Thermophoresis experiment were performed on Monolith NT.115 provided by NanoTemper Technologies Rus. We thank Mr. Edward Ramsay for the help in writing the manuscript.

